# IGF1 Signaling in Temporomandibular Joint Fibrocartilage Stem Cells Regulates Cartilage Growth and Homeostasis in Mice

**DOI:** 10.1101/2021.11.19.469277

**Authors:** Ruiye Bi, Xueting Luo, Qianli Li, Peiran Li, Yi Fan, Binbin Ying, Songsong Zhu

## Abstract

**Objective:** Investigate functional roles of *Igf1* in fibrocartilage stem cell (FCSC) for temporomandibular joint (TMJ) cartilage growth and homeostasis.

**Methods:** *Gli1-CreER*^*+*^; *Rosa*^*TdTomato*^ mice were used for validating FCSCs lineage labeling efficiency. In *Gli1-/Col2-CreER*^*+*^; *Igf1*^*fl/fl*^ mice, TMJ cartilage morphological and functional changes were characterized at 4 weeks and 5 months after *Igf1* deletion. H&E, Safranine O and immuno-histochemistry staining were performed. FCSCs specificity were characterized using EdU and TUNEL staining. A unilateral anterior crossbite (UAC) mouse model was generated for mimicking TMJ osteoarthritis status.

**Results:** In *Gli1-CreER*^*+*^; *Rosa*^*TdTomato*^ mice, RFP labeled FCSCs showed favorable proliferative capacity. 4 weeks after *Igf1* deletion, Gli1^+^ and Col2^+^ cell lineages led to distinct pathological changes of TMJ cartilage morphology. A more serious reduction of cartilage thickness and cell density were found in the superficial layers in *Gli1-CreER*^*+*^; *Igf1*^*fl/fl*^ mice. 5 months after *Igf1* deletion, more severe disordered cell arrangement in TMJ cartilage were found in both groups with Gli1^+^ and Col2^+^ specific deletion of *Igf1*. Immunostaining showed that PI3K/Akt signaling pathway was blocked in the superficial layers of TMJ in *Gli1-CreER*^*+*^; *Rosa*^*TdTomato*^ mice. Finally, deletion of *Igf1* in FCSCs significantly aggravated osteoarthritis (OA) phenotypic changes in TMJ in UAC mice model, characterized in decreased cartilage thickness, cell numbers and loss of extracellular matrix secretion.

**Conclusion:** *Igf1* deletion disrupted stem cell functions of FCSCs, leading to disordered cell distribution during TMJ growth, as well as exaggerated the OA process in TMJ under pathological condition. In TMJ cartilage, *Igf1* expression in FCSCs is critical for PI3K/Akt activation, which may be involved in regulating FCSCs self-renewal and differentiation.

## Introduction

Temporomandibular joint (TMJ) cartilage serves as the center of greatest growth of mandibular skeleton system. During embryonic development, extension of the jaw is controlled by proliferation of chondrocytes within Meckel’s cartilage, which acts as a scaffold for mandibular skeletal morphogenesis (1). This process requires orchestrated interactions between various cell types in TMJ. In TMJ fibrocartilage, it consists of various proportions of both fibrous and cartilaginous tissue. The coated fibrocartilage, instead of hyaline cartilage, showed distinct physiological and pathological process during TMJ homeostasis and injury from cartilage in other joints. Once TMJ fibrocartilage is damaged, it can cause permanent tissue loss and disability, including limitation of mandibular function and growth(2).

Due to restricted self-healing capacity in condylar cartilage, traditional clinical treatment strategies have limited symptom-modifying and structure-modifying effects to restore impaired TMJ cartilage. In previous studies, growth factors have been demonstrated to influence proteoglycan production and deposition. Insulin-like growth factor 1 (IGF1), as one of these growth factors, showed anabolic effect on promoting cell survival and proliferation of chondrocytes, as well as biosynthesis of cartilage matrix macromolecules (3-5). *In vitro*, IGF1 has a stimulating effect for proteoglycan production in a dose-dependent manner (6), and a combination of cell intrinsic and paracrine signals regulate chondrocyte proliferation, which remarkably improved collagen synthesis and mechanical properties of newly formed cartilage. On the other hand, IGF1 activation also shows anti-inflammatory effects in cytokine challenged cartilage tissue (3, 7). For these properties, IGF-1 has been with significant interest as a potential treatment target for TMJ cartilage repair.

Recently, studies have revealed that fibrocartilage stem cells (FCSCs) population located in the superficial zone (SZ) of TMJ cartilage in different species(8, 9), which cells were found with critical roles as cell resources for TMJ homoeostasis and repair (9, 10), and was able to interact with endothelial cells in vascularized bone formation(11). In TMJ osteoarthritis (TMJOA), FCSCs cell features were regulated by various signaling pathways, including Wnt, Notch and TNF-α/Nf-κB signaling (9, 12, 13). During TMJ growth, besides the phenomenon that FCSCs can differentiate toward chondrocytes though, how FCSCs cell fate was influenced and regulated during TMJ development remain to be elucidated. In this current study, we chose to investigate the functional role of IGF1 signaling in FCSCs regulation during TMJ growth and homeostasis. To accomplish this purpose, we used the Cre/Lox-P system to generate an inducible FCSCs-specific *Igf*1 deletion mouse model (*Gli1-CreER*^*+*^; *Igf1*^*fl/fl*^), as well as a chondrocyte-specific *Igf1* deletion mouse (*Col2-CreER*^*+*^; *Igf1*^*fl/fl*^). In our study, we found that both *Gli1-CreER*^*+*^; *Igf1*^*fl/fl*^ mice and *Col2-CreER*^*+*^; *Igf1*^*fl/fl*^ mice manifested with postnatal TMJ cartilage growth restriction. However, the histological changes of TMJ cartilage in these two models were distinct from each other, suggesting that IGF1 signaling activation modulates both FCSCs and chondrocytes in TMJ. Interestingly, deletion of IGF1 signal in FCSCs impaired PI3K-Akt transduction, which was consistent with restrained cell proliferation in the superficial and proliferative zones of TMJ cartilage, suggesting a crucial role for FCSCs-derived IGF1 activation in determination of TMJ cartilage cell proliferation and apoptosis. Finally, we showed that deletion of IGF1 in FCSCs would exaggerate the OA-like cartilage damage in TMJ in a unilateral anterior crossbite (UAC) mouse model.

## Materials and Methods

### Ethical approval information

This study was performed in strict accordance with the recommendations in the Guide for the Care and Use of Laboratory Animals at Sichuan university. Animal procedures were performed according to protocols approved by Animal Ethics Committee at Sichuan university (WCHSIRB-D-2020-431). All animal experiments were followed the ARRIVE (Animal Research: Reporting of In Vivo Experiments) guidelines.

### Animals

Gli1^tm3(creERT2)Alj^/J mice (*Gli1-CreER*^*T2*^, JAX#007913), B6.Cg-Gt(ROSA)26Sor^tm14(CAG-tdTomato)Hze^/J mice (*Tm*^*fl/fl*^, JAX#007908), *Igf1*^*tm1Dlr*^*/J* mice (*Igf1*^*fl/fl*^, JAX#016831) and Tg(Col2a1-cre/ERT)KA3Smac/J (*Col2-CreER*^*T*^,JAX#006774) were obtained from Jackson Laboratory. To generate *Igf1*-conditional knockout mice, *Igf1*^*fl/fl*^ mice were crossed with *Gli1-CreER*^*T2*^ mice and *Col2-CreER*^*T*^ mice respectively.

### Power calculation

The number of mice per experimental group was based on a power calculation using data from our preliminary experiments. We found that a sample size of 6 mice per group would provide greater than 80% power to detect at least a 50% difference between groups in RFP^+^ cell numbers in *Gli1-CreERT*^*+*^; *Igf1*^*fl/fl*^; *Tm*^*fl/fl*^ mice. To account for the possibility of loss of mice due to death before study completion, we used an N of 9 for RFP^+^ cell number counting. The same power calculation strategies were used in *Col2-CreERT*^*+*^; *Igf1*^*fl/fl*^; *Tm*^*fl/fl*^ mice.

### Tamoxifen administration

In the TMJ cartilage growth experiment, *Gli1-CreER*^*+*^; *Igf1*^*fl/fl*^ mice and *Col2-CreER*^*+*^; *Igf1*^*fl/fl*^ mice were injected intraperitoneally with tamoxifen at P12 (75mg/kg body weight/day, for 4 consecutive days). 4 weeks and 5 months after the first injection, mice were sacrificed for histologic analysis. The Cre-negative littermates were used as controls. In the unilateral anterior crossbite (UAC) animal model experiment, *Gli1-CreER*^*+*^; *Igf1*^*fl/fl*^ mice and *Col2-CreER*^*+*^; *Igf1*^*fl/fl*^ mice were injected intraperitoneally with tamoxifen at the age of 6 weeks (75mg/kg/day, for 4 consecutive days). 4 weeks after the first injection, mice were sacrificed for further analyses. The Cre-negative littermates were used as controls.

### Unilateral anterior crossbite (UAC) mouse model generation

2 days after the first Tamoxifen injection, g*li1-CreER*^*+*^; *Igf1*^*fl/fl*^ mice and *Igf1*^*fl/fl*^ mice were anaesthetized by 70 μg/g Ketamine and 15 μg/g Xylazine intraperitoneally. *Igf1*^*fl/fl*^ mice equivalent in age and weight were randomly divided into the Sham group and UAC group. Low speed electric drill was used to cut needles from 20ml syringe into 5-8mm long metal tube. A curved metal tube was bonded to the left mandibular incisor of the mouse by zinc phosphate cement. The metal tubes were checked every day in case they fell off. Only those mice whose metal tubes remained in place during the entire observation period were included in this experiment.

### Label-retaining cells

0.1mg of 5-ethynyl-2’-deoxyuridine (EdU) (Invitrogen, A10044) dissolved in PBS (50ul) was administered to 3-week *Gli1-CreER*^*+*^; *Igf1*^*fl/fl*^ mice and *Col2-CreER*^*+*^; *Igf1*^*fl/fl*^ mice IP four hours before sacrificed. The Click-iT Imaging Kit (Invitrogen, C10338) was used to detect EdU-positive cells on paraffin embedded sections by immunofluorescence. EdU labled^+^ cells were counted by an independent researcher

### TUNEL assay

TUNEL staining including antigen retrieval (microwave 5min x 2) and staining process was performed following manufacturer’s instruction (11864795910, Roche Applied Science). FITC-labeled TUNEL-positive cells were imaged under fluorescent microscopy. DAPI was used for counterstaining.

### Histology and Immunohistochemistry

Tissue samples were fixed in 4% paraformaldehyde for 24 hours. Mice condyles were decalcified in 15% ethylenediaminetetraacetic acid (Sigma Aldrich) for 4 weeks. Samples were embedded in paraffin and 5μm sections were cut. Serial tissue sections were stained with hematoxylin and eosin (H&E) and Safranine O. For immunofluorescent staining, sections were blocked with 10% goat serum for 1 hour at room temperature and incubated with primary antibody overnight at 4°C. Sections were then stained at room temperature for 1 hour with second antibody. DAPI (Vector Laboratories) was used for nuclear counterstaining. The primary antibodies used in our experiments were mouse anti-RFP (1:100, sc-390909, Santa Cruz), mouse anti-Col2a1 (1:100, sc518040, Santa Cruz), rabbit anti-SOX9 (1:100, Abcam), rabbit anti P-Akt (1:100,9271, Cell Signal Technology), MCM2(1:100,3619, Cell Signal Technology). The second antibodies used in our experiments were Alexa Fluor 568 goat anti mouse IgG antibody (1:500, A-11004, Invitrogen), Alexa Fluor 568 goat anti rabbit IgG antibody (1:500, A-11036, Invitrogen), Alexa Fluor 488 goat anti rabbit IgG antibody (1:500, A-11034, Invitrogen) and Alexa Fluor 488 goat anti mouse IgG antibody (1:500, A32723, Invitrogen). For immunochemistry, sections were blocked with 10% rabbit serum for 15min at room temperature and then incubated with goat anti mouse IGF1(1:40, AF791, R&D System) overnight at 4°C. The reaction was interrupted by biotin-labeled IgG (PV-9003, Zhongshan Golden Bridge) and an avidin–peroxidase complex at 37 °C for 30 min each. For the 3,3-diaminobenzidine (DAB) staining step, the slides were colored for 30s and then the nucleus was counterstained with hematoxylin for 30s. For all histological semi-quantification analyses, each section was blindingly counted by an independent researcher for twice, and the average number counted was calculated as the final counted number.

### Statistical Analysis

Statistical testing was performed on results with no animals or raw data points excluded. The staining results were analyzed by Image J 1.51 (Leeds precision instruments, USA). All statistics were calculated using Prism8 (GraphPad Software). The statistical significance between 2 groups was determined using paired Student’s t-test, and the statistical significance between 3 groups was determined using One-way ANOVA with multiple comparisons. The difference was statistically significant when p value< 0.05.

## Results

### Generation and validation of *Igf1* deletion in Gli1^+^ FCSCs lineage and Col2^+^ chondrocyte lineage

Gli1, an essential hedgehog signaling transcription factor, functions in undifferentiated cells during embryogenesis. Gli1 was also identified as one of the markers for mesenchymal-derived stem cells in the craniofacial system, including cranial suture (14), incisor (15), peridontium(16) and alveolar bone marrow(17). To investigate whether Gli1 serves as a specific marker for FCSCs, we investigated Gli1 expression pattern in the TMJ cartilage development. We bred *Gli1-CreER*^*+*^ mice with *Rosa*^*Tdtomato*^ reporter mice, whose tissue expressed fluorescence TdTomato in the absence of Cre-mediated recombination (Figure 1A). 5 days after tamoxifen injection, sections of the TMJ showed scattered expression of red fluorescent protein (RFP) along the surface of the fibrocartilage zone in mice (Figure 1B). 15 days after tamoxifen injection, robust RFP^+^ cells were observed distributed not only in the SZ, but also in the polymorphic zone (PZ), suggesting that Gli1^+^ cell lineage proliferated and migrated from SZ to deeper TMJ cartilage zones. Next, to generate *Igf1* specific deletion in FCSCs in mice, Gli1-Cre^+^ mice were bred to *Igf1* floxed mice (*Igf1*^*fl/fl*^) and resultant *Gli1-CreER*^*+*^; *Igf1*^*fl/+*^ mice were crossed to *Igf1*^*fl/fl*^ mice to generate KO mice (*Gli1-CreER*^*+*^; *Igf1*^*fl/fl*^) and littermate control mice (*Igf1*^*fl/fl*^) (Figure 1D). To validate FCSCs-specific deficiency of IGF1 in our *Gli1-CreER*^*+*^; *Igf1*^*fl/fl*^ mice, we measured *Igf1* mRNA expression levels in TMJ cartilage tissues from *Gli1-CreER*^*+*^; *Igf1*^*fl/fl*^ mice and *Igf1*^*fl/fl*^ mice. *Igf1* mRNA expression was reduced by 81% in cartilage cells from *Gli1-CreER*^*+*^; *Igf1*^*fl/fl*^ mice (Figure 1E). Importantly, *Igf1* mRNA expression was unchanged in *Igf1*^*fl/fl*^ mice. There have been previous reports of Gli1-Cre mediating germline gene deletion of certain floxed alleles; however, this did not occur in *Gli1-CreER*^*+*^; *Igf1*^*fl/fl*^ mice. Meanwhile, we performed immuno-histochemical staining of IGF1 protein expressions in TMJ cartilage. A significantly reduction of IGF1 protein expression was found in *Gli1-CreER*^*+*^; *Igf1*^*fl/fl*^ mice in TMJ cartilage SZ and PZ area (IGF1^+^ cells% from (26.3±7.3)% to (6.6±3.6)%), which signified the success of conditional deletion of *Igf1* gene expression in FCSCs (Figure 1 G).

**Figure 1.**
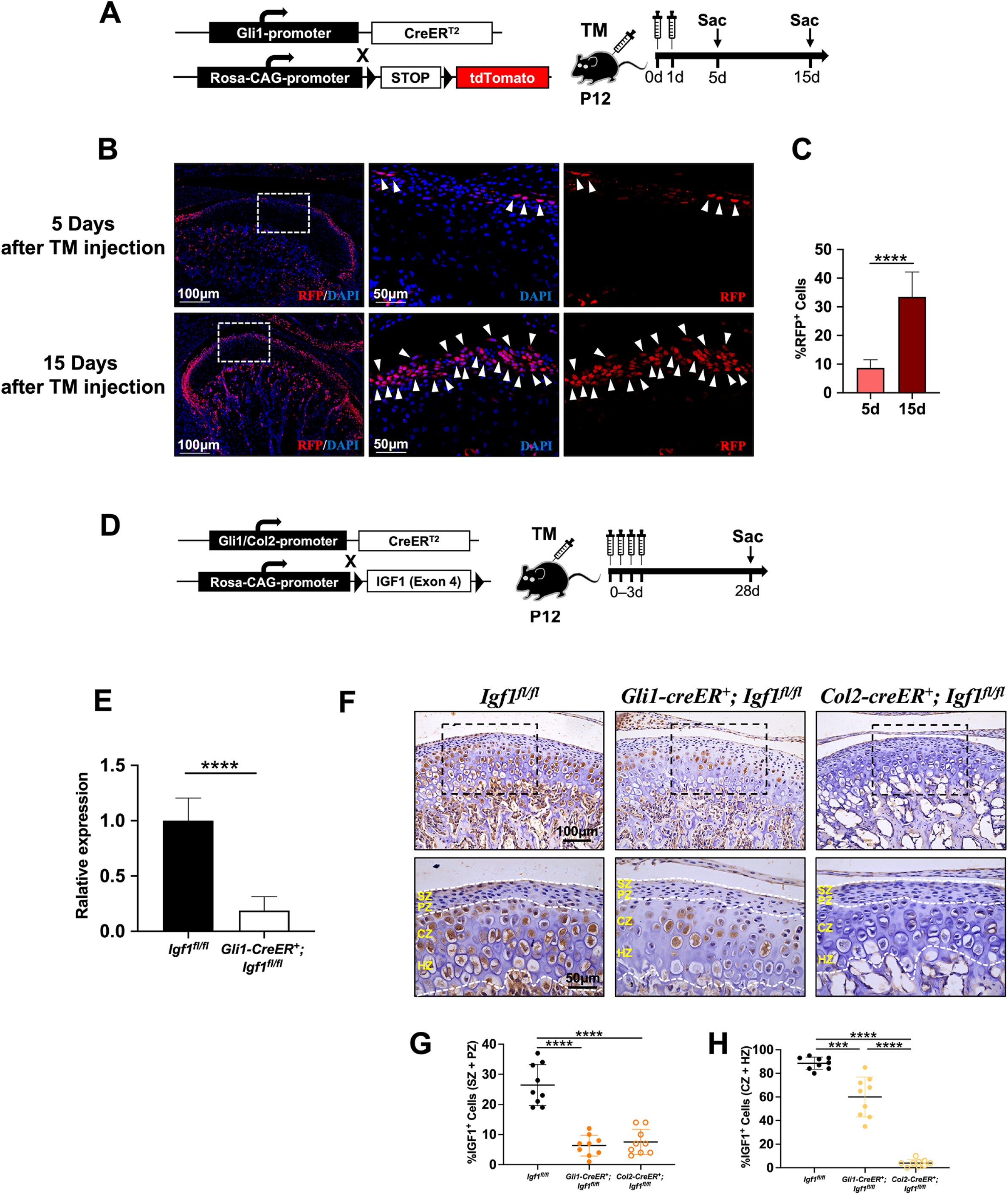
Generation and validation of *Igf1* deletion in Gli1^+^ FCSCs *in vivo*. (A) Mating tactics for generating *Gli1-CreER; Rosa*^*TdTomato*^ mice. (B) Representative images of RFP^+^ cell distributions in TMJ cartilage, 5 and 15 days after tamoxifen injection, RFP activation was in TMJ sections. (C) RFP^+^ Cell numbers in TMJ cartilage 15 and 15 days after Tm injection. (D) Mating tactics for generating *Gli1-CreER*^*2*^; *Igf1*^*fl/fl*^ mice and *Col2-CreER; Igf1*^*fl/fl*^ mice. (E) mRNA expression and (F) protein expression of IGF1 after specific deletion of IGF1 in *Gli1-CreER*^*2*^; *Igf1*^*fl/fl*^ mice and *Col2-CreER; Igf1*^*fl/fl*^ mice. IGF1KO efficiency in (G) SZ and PZ and (H) CZ and HZ. N=9, ***p<0.001, ****p<0.0001. SZ: superficial zone; PZ: proliferative zone; CZ: chondrogenic zone; HZ: hypertrophic zone.

To compare the different regulating role of *Igf1* in other cells in TMJ, we also generated a chondrocyte specific *Igf1* deletion mouse model. *Col2-CreER*^*+*^ mice were bred to *Igf1*^*fl/fl*^ mice and resultant *Col2-CreER*^*+*^; *Igf1*^*fl/+*^ mice were crossed to *Igf1*^*fl/fl*^ mice to generate KO mice (*Col2-CreER*^*+*^; *Igf1*^*fl/fl*^) and littermate control mice (*Igf1*^*fl/fl*^). To validate Col2-specific deficiency of IGF1 in our *Col2-CreER*^*+*^; *Igf1*^*fl/fl*^ mouse, immuno-histochemical staining of TMJ cartilage showed a marked reduction of IGF1 protein expression in the chondrogenic zone (SZ) and hypertrophic zone (HZ) of TMJ cartilage, (IGF1^+^ cells% from (88.5±5.1)% to (4.0±3.0)%) (Figure 1H).

#### Deletion of Igf1 in Gli1^+^ cell lineage and Col2^+^ cell lineage leads to distinct pathological changes of TMJ cartilage morphology

We next investigated the role of IGF1 in zonal organization of condylar articular cartilage in juvenile and adult mice. *Gli1-CreER*^*+*^; *Igf1*^*fl/fl*^ mice, *Col2-CreER*^*+*^; *Igf1*^*fl/fl*^ mice and *Igf1*^*fl/fl*^ mice were injected with tamoxifen at P12-P15, and mice were harvested after 4 weeks and 5 months (Figure 2A, S1A). In *Igf1*^*fl/fl*^ mice, the TMJ cartilage displayed well-organized multi-layer cell distribution with distinct four-layer cartilaginous zones. In *Gli1-CreER*^*+*^; *Igf1*^*fl/fl*^ mice and *Col2-CreER*^*+*^; *Igf1*^*fl/fl*^ mice group, their TMJ cartilage layers were found with disordered arrangement (Figure 2B). Semi-quantification of cell numbers and cartilage thickness showed that the cartilage thickness in both *Gli1-CreER*^*+*^; *Igf1*^*fl/fl*^ mice and *Col2-CreER*^*+*^; *Igf1*^*fl/fl*^ mice were not significantly decreased, while the superficial zone (SZ) and proliferative zone (PZ) had a trend of decreased thickness compared with control group with no statistical significance (Figure 2C-D). Cell density in TMJ cartilage were found significantly decreased in both *Gli1-CreER*^*+*^; *Igf1*^*fl/fl*^ mice (by 39.8%) and *Col2-CreER*^*+*^; *Igf1*^*fl/fl*^ mice (by 34.9%) (Figure 2E). Especially when we compared cell numbers in the SZ and PZ, the cell density of *Gli1-CreER*^*+*^; *Igf1*^*fl/fl*^ mice group were dramatically reduced compared with the other groups, which were only 66.9% of the *Col2-CreER*^*+*^; *Igf1*^*fl/fl*^ mice and 80.5% of the control mice (Figure 2F). We further performed Safranine O staining, detecting a slight increase of the proteoglycan in cartilage in *Col2-CreER*^*+*^; *Igf1*^*fl/fl*^ mice, which phenomenon suggested the initiation of cartilage degradation (Figure 2G-H). Consistently, we also found a significantly decrease of collagen II expression in both *Gli1-CreER*^*+*^; *Igf1*^*fl/fl*^ mice and *Col2-CreER*^*+*^; *Igf1*^*fl/fl*^ mice (Figure 2I-J).

**Figure 2.**
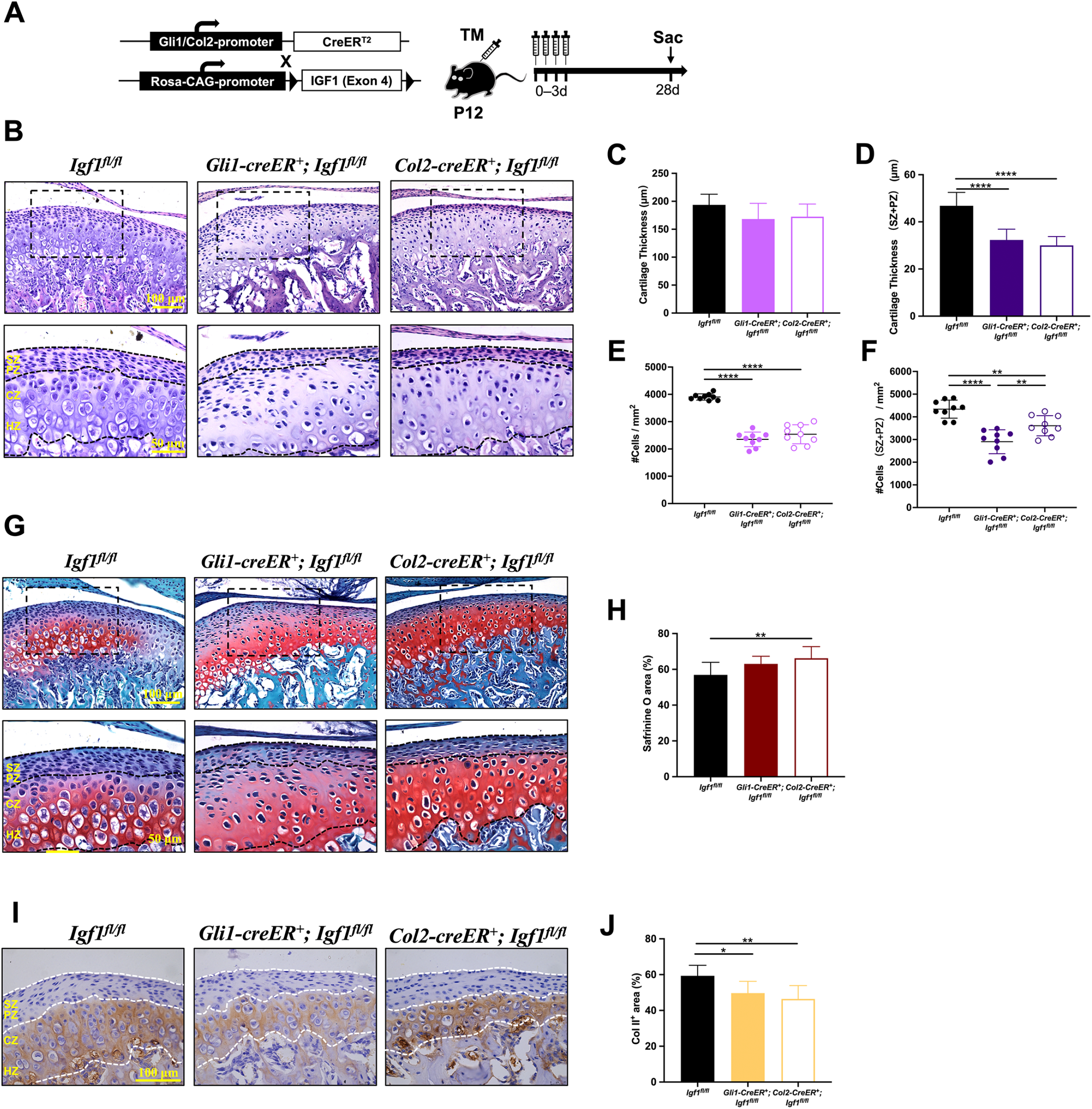
Deletion of *Igf1* in Gli1^+^ cell lineage and Col2^+^ cell lineage. (A) Mating tactics for generating *Gli1-CreER; Igf1*^*fl/fl*^ mice and *Col2-CreER; Igf1*^*fl/fl*^ mice. (B) H&E staining of TMJ cartilage at 4 weeks after Tm injection, and (C) cartilage thickness, (D) cartilage thickness in SZ and PZ, (E) cell density, and (F) cell density in SZ and PZ were quantified. (G) Safranine O staining of TMJ cartilage at 4 weeks after Tm injection, and (H) Safranine O^+^ area was quantified. (I) Col II immune-histochemical staining of TMJ cartilage at 4 weeks after Tm injection, and (J) Col II^+^ area was quantified. N=9. *p<0.05, **p<0.01, ***p<0.001, ****p<0.0001. SZ: superficial zone; PZ: proliferative zone; CZ: chondrogenic zone; HZ: hypertrophic zone.

At the same time, when we observed cartilage morphological changes in *Col2-CreER*^*+*^; *Igf1*^*fl/fl*^ mice, we found that besides decreased cell numbers in chondrogenic zones, the cell numbers were not significantly affected in the SZ area (Figure 2E-H), The cell arrangement of SZ in *Col2-CreER*^*+*^; *Igf1*^*fl/fl*^ mice were preserved with disordered cell distribution in the inferior chondrocytes (Figure 2E-F).

#### Deletion of Igf1 leads to long-term disruption in growth and differentiation in both Gli1d^+^ cell lineage and Col2^+^ cell lineage

5 months after *Igf1* deletion in FCSCs (Figure S1A), a widespread cartilage disarrangement occurred in *Gli1-CreER*^*+*^; *Igf1*^*fl/fl*^ mice (Figure S1B). The original multi-layer cell distribution was replaced by an osteoarthritis (OA)-like morphological changes, including dramatically decrease of cell numbers in all layers, rough surface in the SZ area and clustering of the chondrocytes (Figure S1B, C-F). Proteoglycans were not observed in chondrogenic zone of the *Gli1-CreER*^*+*^; *Igf1*^*fl/fl*^ mice, signifying the loss of function of chondrocytes in matrix secretion at this stage (Figure S1G-H).

Interestingly, the TMJ cartilage in *Col2-CreER*^*+*^; *Igf1*^*fl/fl*^ mice showed significantly different morphological changes at this stage compared with 4 weeks after *Igf1* deletion. The SZ area became no-longer intact, with significant loss of cell numbers and disordered cell distributions (Figure S1B, C-F). We also found that the proteoglycan secretion of chondrocytes was moderately decreased at this stage (Figure S1G,-H). This phenomenon hint that the differentiation capacity of FCSCs toward cartilage was prostrated, which could result in interrupted TMJ cartilage development.

#### Deletion of Igf1 leads to disruption of chondrogenic capacity of FCSCs in vivo

Since *Igf1* deletion markedly affected the self-renewal and chondrogenic ability, we further investigated the feature changes of resident pluripotent cells in the TMJ cartilage surface. We observed statistically increased cell apoptosis in the TMJ SZ and PZ area of *Gli1-CreER*^*+*^*;Igf1*^*fl/fl*^ mice from (4.7±1.9)% to (10.9±5.1)% (Figure 3A-B). Instead, the cell proliferation capacity in the SZ and PZ cells were significantly decreased from (14.4± 3.4)% to (3.5±1.3)% by EDU staining (Figure 3C-D). To investigate the cell fate change of FCSCs, we performed immunofluorescent staining of SOX9, and analyzing changes of chondrogenic capacity of cells in SZ and PZ. In *Gli1-CreER*^*+*^*;Igf1*^*fl/fl*^ mice, the percentage of SOX9^+^ cells in the SZ and PZ were dramatically decreased, from (15.7±4.9%) to (1.5 ±1.7%)(Figure 3 E-F).

**Figure 3.**
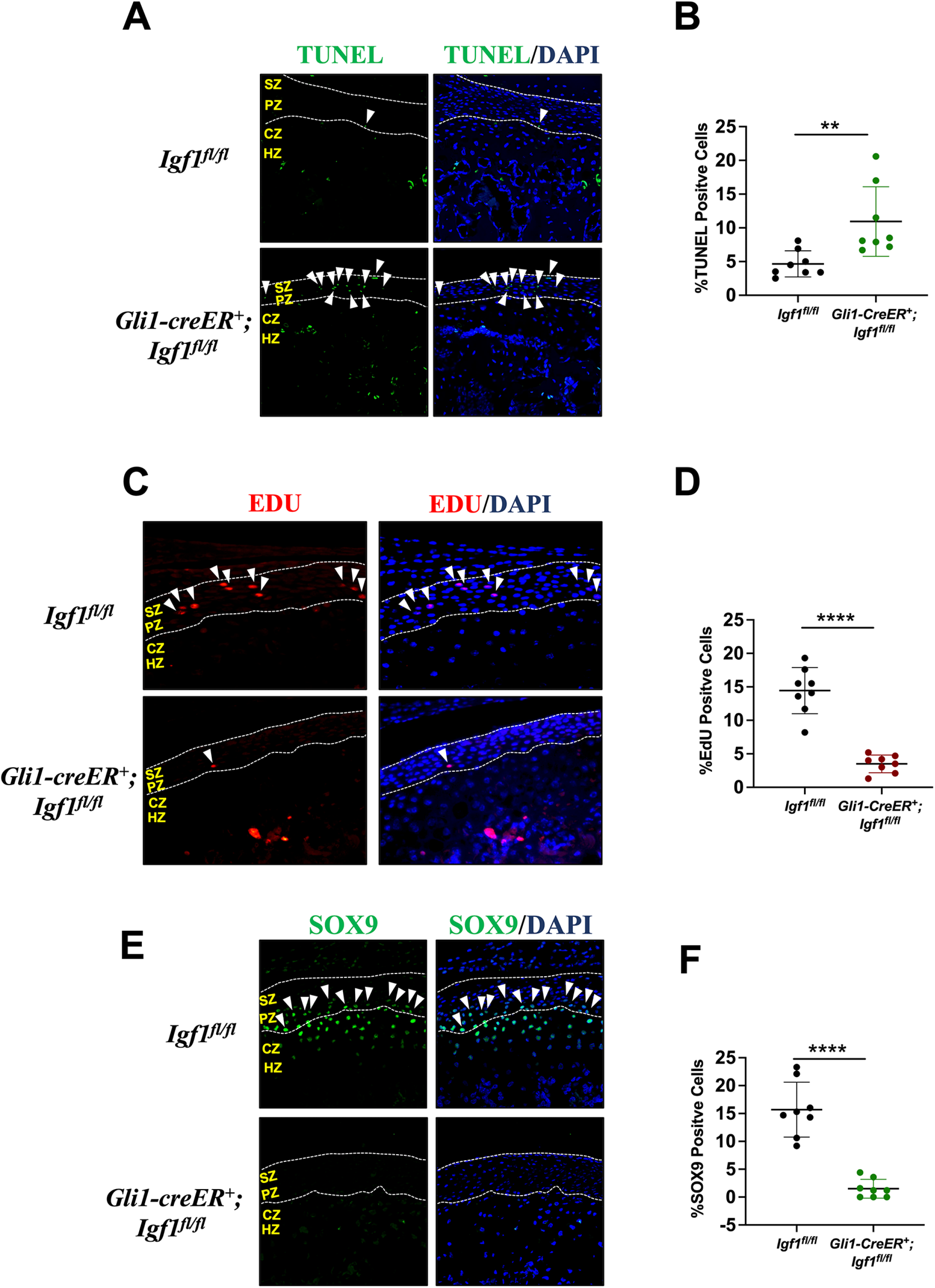
Deletion of *Igf1* leads to disruption of chondrogenic capacity of FCSCs in vivo. (A) TUNEL staining of TMJ cartilage in *Gli1-CreER; Igf1*^*fl/fl*^ mice, and (B) TUNEL^+^ cells were quantified. (C) EdU staining of TMJ cartilage in *Gli1-CreER; Igf1*^*fl/fl*^ mice, and (D) EdU^+^ cells were quantified. (E) SOZ9 immunostaining of TMJ cartilage in *Gli1-CreER; Igf1*^*fl/fl*^ mice, and (F) SOX9^+^ cells were quantified. **p<0.01, ****p<0.0001. SZ: superficial zone; PZ: proliferative zone; CZ: chondrogenic zone; HZ: hypertrophic zone.

To illuminate the mechanistic role of IGF1 signaling in FCSCs proliferation and apoptosis, we performed immunofluorescent staining of p-AKT, as well as MCM, both of which are key modulators in cartilage cell renewal, differentiation, and apoptosis. We found that under physiological conditions, p-AKT and MCM2 were mainly activated in the SZ and PZ (Figure 4A). When IGF1 signaling was blocked in FCSCs (*Gli1-CreER*^*+*^; *Igf1*^*fl/fl*^), both p-AKT and MCM2 expressions were found significantly reduced in SZ and PZ (Figure 4 A-D). Taken together, these data suggested that *Igf1* gene expression was involved in the differentiation and apoptosis regulation of condylar cartilage by FCSCs, which signaling may play its roles in maintaining TMJ cartilage development and homeostasis *via* PI3K-AKT signaling pathway.

**Figure 4.**
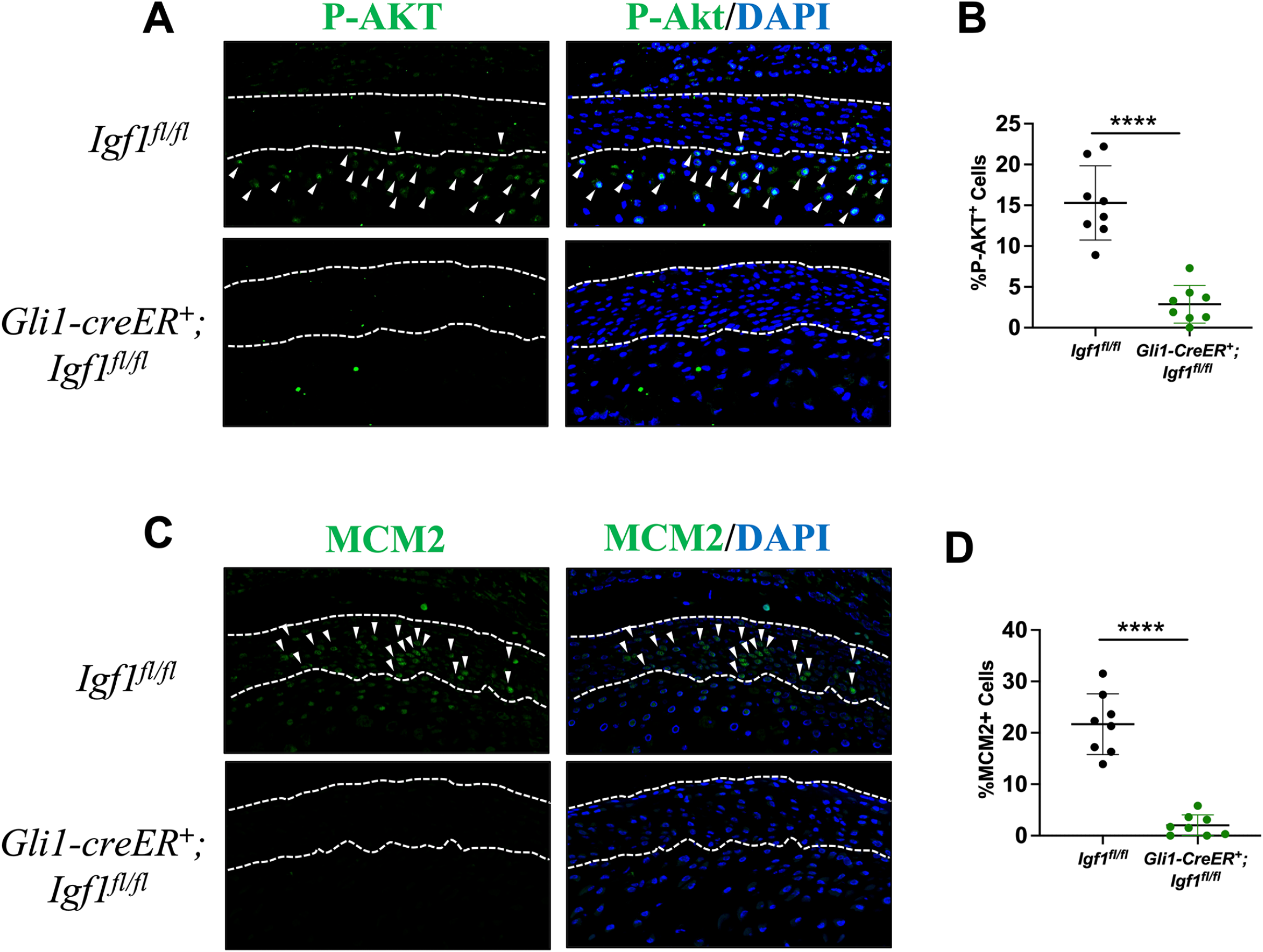
Deletion of *Igf1* leads to PI3K/Akt signaling inhibition in TMJ surface. (A) P-Akt immunostaining of TMJ cartilage in *Gli1-CreER; Igf1*^*fl/fl*^ mice, and (B) P-Akt^+^ cells were quantified. (C) MCM2 immunostaining of TMJ cartilage in *Gli1-CreER; Igf1*^*fl/fl*^ mice, and (B) MCM2^+^ cells were quantified. ****p<0.0001.

#### Deletion of Igf1 aggravated OA phenotype in TMJ cartilage in UAC mice model

After finding that *Igf1* gene deletion led to disordered arrangement of TMJ cartilage during development, we continued to explore whether *Igf1* gene were involved in the process of TMJ pathological changes. We generated a unilateral anterior crossbite (UAC) mouse model, which model were shown with a slight OA phenotype (Figure 5A). 4 weeks after model generation, the *Igf1*^*fl/fl*^ mice with UAC displayed moderately decreased cartilage thickness, with cell density decrease from (4172±301)/mm^2^ to (3371±325)/mm^2^ (Figure 5E-F) with slightly disarranged multilayered cell distribution (Figure 5B). Cartilage thickness of PZ and CZ were thinner than control group, and the cell density were also found decreased after model generation (Figure 5C-D). At the same time, proteoglycan secretion was found increased (Figure 5 G-H). These morphological changes represented an early OA change in the TMJ. Interestingly, when *Igf1* was deleted in FCSCs, UAC model generation led to a more serious OA phenotypic change in TMJ cartilage. The cartilage thickness was dramatically decreased with loss of layered cell distribution in the TMJ, as well as marked cell number decrease (from (3371±325)/mm^2^ to (2875±241)/mm^2^) (Figure 5B-F). In the chondrogenic zone, chondrocytes lost their capacity of matrix secretion, showing little positive signal with the Safranine O staining, as well as Col II immunostaining (Figure 5G-H, S2B-C). Which phenomenon represent the late stage of OA like changes in these mice.

**Figure 5.**
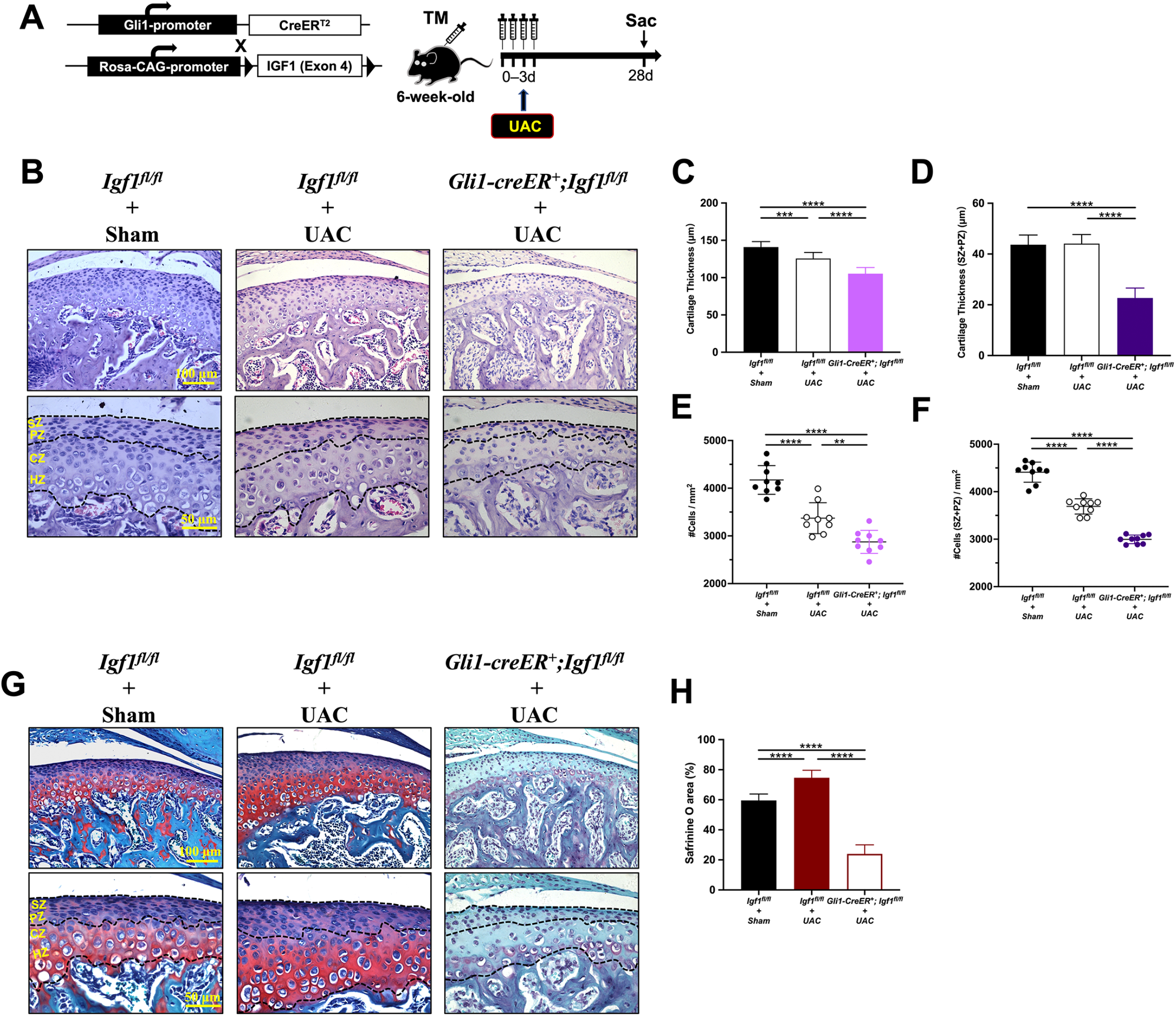
Deletion of *Igf1* aggravated OA phenotype in TMJ cartilage in UAC mice model. (A) Strategies for generating the *Gli1-CreER; Igf1*^*fl/fl*^ UAC mice mouse model. (B) H&E staining of TMJ cartilage at 4 weeks after Tm injection, and (C) cartilage thickness, (D) cartilage thicknes in SZ and PZ, (E) cell density, and (F) cell density in SZ and PZ were quantified. (G) Safranine O staining of TMJ cartilage at 4 weeks after Tm injection, and (H) Safranine O^+^ area was quantified. N=8-9, *p<0.05, **p<0.01, ***p<0.001.

## Discussion

In previous studies, IGF1 was proved to be a ubiquitously expressed hormone and plays an essential role in postnatal growth(18, 19), and approximately 75% of circulating IGF1 is derived from the liver(20, 21). In the skeletal system, targeted deletion of IGF1 in chondrocytes(22), osteoblasts(23), or osteocytes(24) produces unique effects on homeostasis and growth in the skeletal system, without affecting circulating IGF concentrations. Previous studies have also hinted a potential role for IGF1 in cartilage biology, especially in chondrocyte proliferation, differentiation and hypotraphy(25-27). However, the studies linking IGF1-mediated signaling with cartilage growth and homeostasis are limited by the fact that many of the effects seen may be due to indirect effects on non-chondrocyte cells.

In the SZ of TMJ cartilage, FCSCs were identified as endogenic resources for cartilage formation, which cells contributed to the differentiation of chondrocytes in juvenile mice(9). However, the key regulator for mediating FCSCs stem cell features are yet to be clarified. In our study, we investigated the functional role of *Igf1* expression in FCSCs using Cre/Lox-P–mediated gene deletion. Although *Gli1-CreER*^*+*^; *Igf1*^*fl/fl*^ mice demonstrated no significant postnatal growth restriction, we did identify cartilage morphological alterations in TMJ, suggesting that *Igf1* is essential for the canonical TMJ cartilage function in chondrogenic stem cell niche.

We provided evidence that *Igf1* in FCSCs and chondrocytes has distinct effect on morphological changes in the TMJ cartilage. At early stage, IGF1 specific deletion in chondrocytes mainly lead to functional disruption of matrix secretion in these cells, while IGF1 ablation in FCSCs lead to milder disruption of cartilage matrix secretion. The extracellular matrix secretion is of great importance for TMJ cartilage quality and strength. These findings showed that autocrine IGF1 in chondrocytes mainly participated in the matrix secretion in TMJ.

At early stage, the proliferation and differentiation capacity of FCSCs in SZ was maintained when IGF1 was specifically deleted in chondrocytes. However, when IGF1 was deleted in FCSCs, it would lead to a more severe cell number loss in the entire cartilage layers, as well as increased apoptosis, decreased proliferation and loss of chondrogenic capacity in SZ and PZ cells. At late stage, deletion of IGF1 signaling in both FCSCs and chondrocyte lead to similar phenotype, showing an OA-like disarrangement in TMJ cartilage. Based on the results from our current study, we find that IGF1 does have an appreciable regulatory role on TMJ cartilage homeostasis, which mainly by affecting the cell proliferation, differentiation and apoptosis capacity of FCSCs, as well as the matrix secretion of chondrocytes.

In conclusion, this study describes a novel role for IGF1 signaling during TMJ cartilage development and homeostasis, which acts as critical modulator of cell differentiation capacities in both FCSCs and chondrocytes. Apart from mediating cell proliferation and differentiation during TMJ cartilage growth, we show that under pathological conditions, *Igf1* deletion appears to be accelerator for cartilage degradation, and may aggravated cartilage damage during TMJOA.

## Acknowledgements

None

## Author contributions

R.B contributed to conception, design, data acquisition, analysis and interpretation, drafted and critically revised the manuscript; X.L contributed to design, data acquisition and analysis, and critically revised the manuscript; Q.L contributed to data acquisition and critically revised the manuscript; P.L contributed to data analysis and critically revised the manuscript; B.Y contributed to design, data analysis and interpretation, and critically revised the manuscript; S.Z contributed to conception, design, data analysis and interpretation, drafted and critically revised the manuscript. All authors gave their final approval and agreed to be accountable for all aspects of the work

---

**Figure S1.**
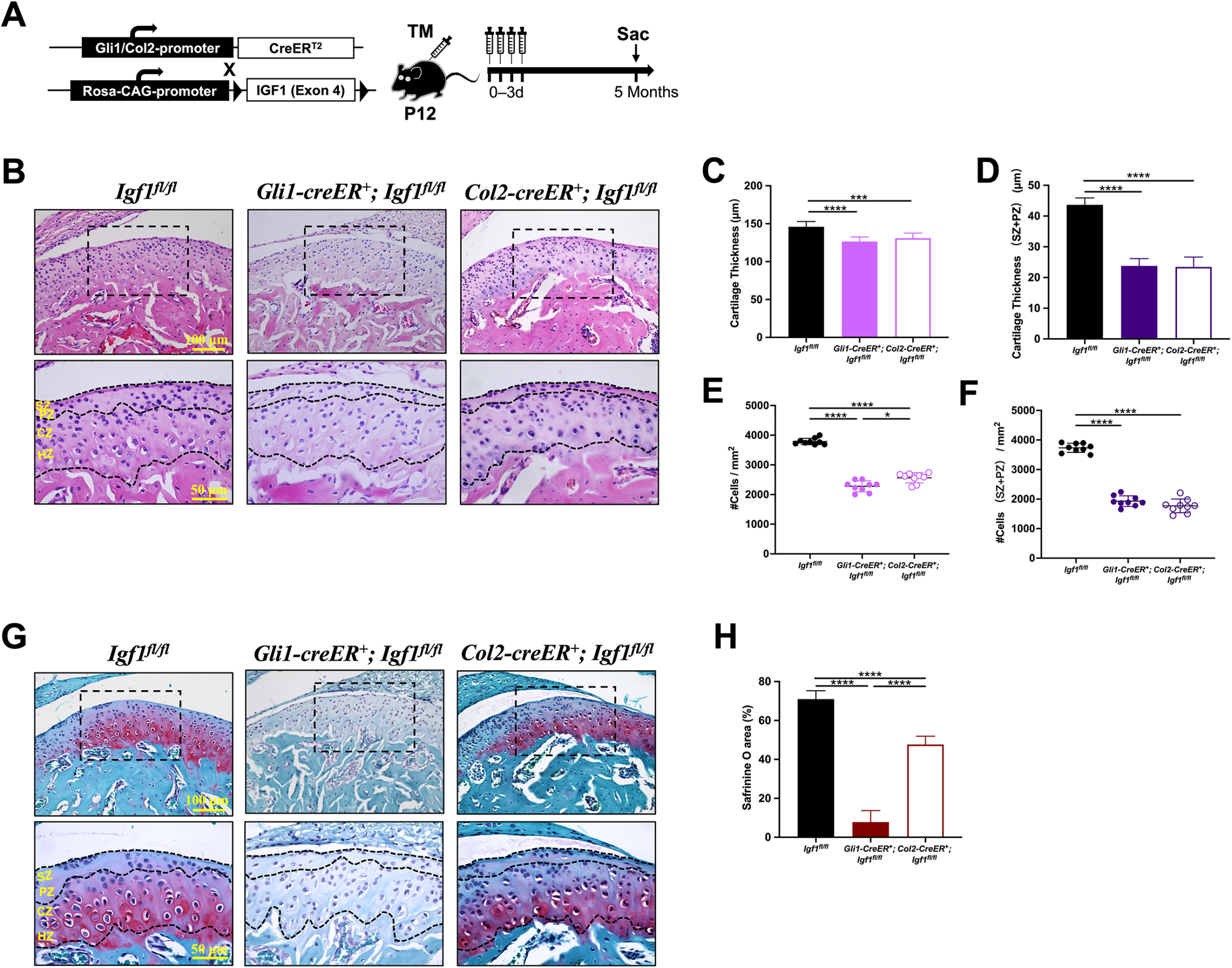
Long-term deletion of *Igf1* in Gli1^+^ cell lineage and Col2^+^ cell lineage. (A) Mating tactics for generating TM inducible long-term deletion of *Igf1* in *Gli1-CreER; Igf1*^*fl/fl*^ mice and *Col2-CreER*^*+*^; *Igf1*^*fl/fl*^ mice. (B) H&E staining of TMJ cartilage at 5 months after Tm injection, and (C) cartilage thickness, (D) cartilage thickness in SZ and PZ, (E) cell density, and (F) cell density in SZ and PZ were quantified. (G) Safranine O staining of TMJ cartilage at 5 months after Tm injection, and (H) Safranine O^+^ area was quantified. N=9. *p<0.05, ***p<0.001, ****p<0.0001. SZ: superficial zone; PZ: proliferative zone; CZ: chondrogenic zone; HZ: hypertrophic zone.

**Figure S2.**
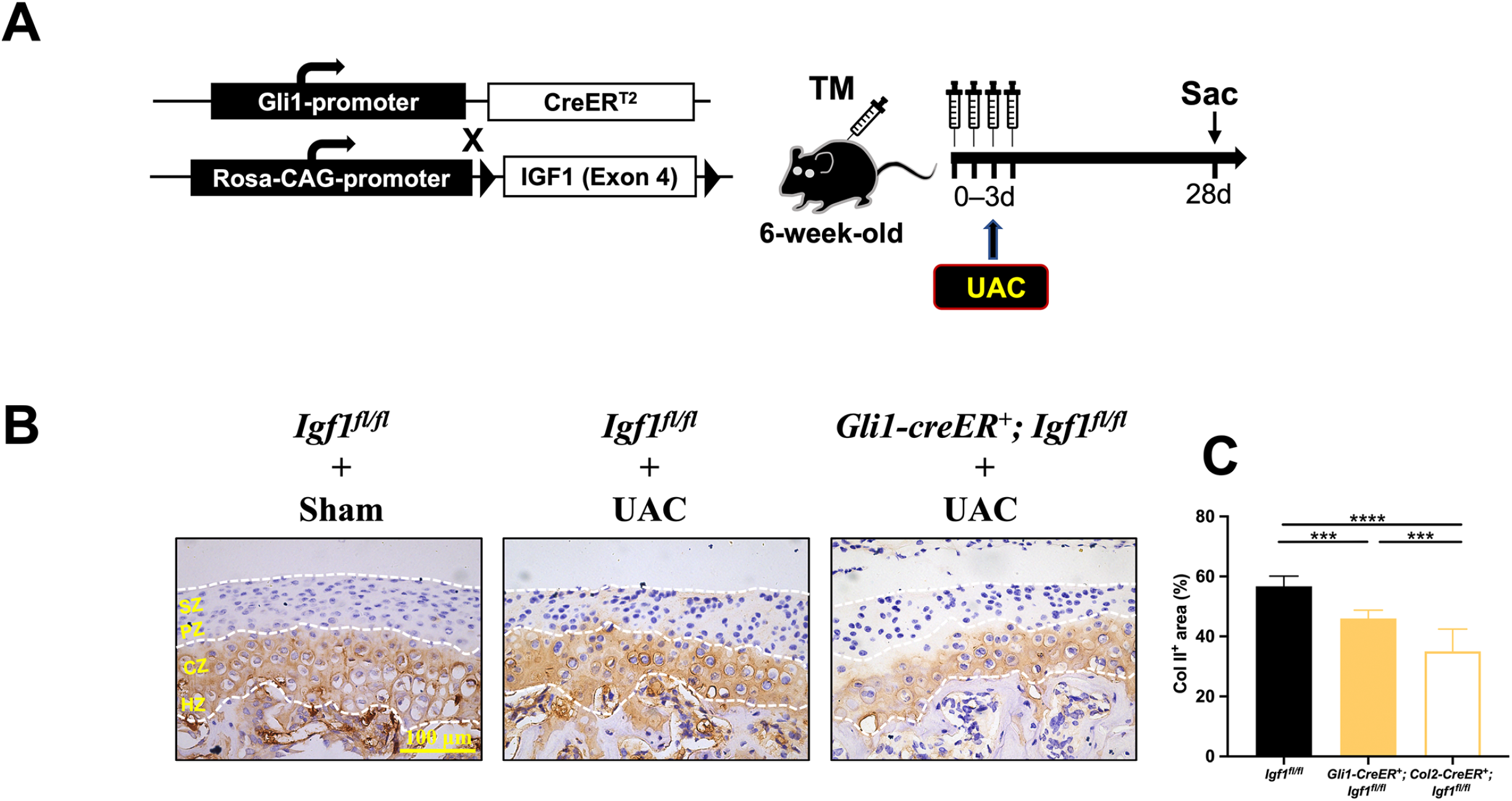
Col Il protein expression changes after *Igf1* deletion in UAC mice model. (A) Strategies for generating the *Gli1-CreER*^*+*^; *Igf1*^*fl/fl*^ UAC mice mouse model. (B) Col II immune-histochemical staining of TMJ cartilage at 4 weeks after Tm injection in UAC mouse model, and (C) Col II^+^ area was quantified. N=9. ***p<0.001, ****p<0.0001. SZ: superficial zone; PZ: proliferative zone; CZ: chondrogenic zone; HZ: hypertrophic zone.

